## Supplementary Methods for "Joint Modeling of Cellular Heterogeneity and Condition Effects with scPCA in Single-Cell RNA-Seq"

### The scPCA model

The scPCA factorisation requires a multi-condition single-cell expression count matrix $\mathbf{X} \in \mathbb{N}_0^{C \times G}$ for $C$ cells and $G$ genes, a design matrix $\mathbf{D} \in \{0,1\}^{C \times Q}$ that encodes a conditioning variable with $Q$ levels, and an optional indicator matrix $\mathbf{B} \in \{0,1\}^{C \times P}$ that may adjust for condition-specific mean gene expression for $P$ groups of cells (usually $P = 1$ or $P = Q$, i.e. scPCA fits a global or condition-specific mean offset(s) respectively) as inputs. We model the elements of the expression count matrix $\mathbf{X} = \{x_{cg}\}$ for cell $c = 1, \ldots, C$ and gene $g = 1, \ldots, G$ as negative binomial distributed, given an unobserved gene expression level $\mu_{cg}$ and a gene-specific over-dispersion $\alpha_g$

$$x_{cg} \sim \mathrm{NB}(\mu_{cg}, \alpha_g). \tag{1}$$

In scPCA, we decompose the unobserved gene expression as a linear function of the conditioning variable encoded in $\mathbf{D}$. Denote the entries of the design and indicator matrix $\mathbf{D} = \{d_{cq}\}$ and $\mathbf{B} = \{b_{cp}\}$ for level $q = 1, \ldots, Q$ and group $p = 1, \ldots, P$ respectively, and let $l_c = \log \sum_{g=1}^{G} x_{cg}$ denote a cell-specific fixed size factor (total number of gene counts). Then,

$$\log \mu_{cg} = \underbrace{l_c}_{\text{Size factor}} + \underbrace{\left( \sum_{p=1}^{P} b_{cp} v_{pg} \right)_{cg}}_{\text{Intercept term}} + \underbrace{\left( \sum_{q=1}^{Q} d_{cq} \cdot \sum_{f=1}^{F} z_{cf} w_{fgq} \right)_{cg}}_{\text{Factor model}} \tag{2}$$

where $w_{fgq}$ represents the entries of a loading weight tensor $\mathbf{W} \in \mathbb{R}^{F \times G \times Q}$ consisting of $f = 1, \ldots, F$ components, $z_{cf}$ the elements a factor weight matrix $\mathbf{Z} \in \mathbb{R}^{C \times F}$ containing the coordinates of each cell with respect to $\mathbf{W}$, and $v_{pg}$ the entries of a mean offset $\mathbf{V} \in \mathbb{R}^{P \times G}$ matrix.

### Model inputs and parameters

#### The expression count matrix

In the following, we will restrict our discussion to a single conditioning variable with levels $q = 1, \ldots, Q$. To jointly model multi-condition single-cell data using scPCA, we require the user to decide on a reference level for comparison. For example, in the common control-treatment setting, the cells of the control condition represent a natural choice. In general, the expression count matrix may take the form

$$\mathbf{X} = \begin{bmatrix} \mathbf{X}^1 \\ \mathbf{X}^2 \\ \mathbf{X}^3 \\ \vdots \\ \mathbf{X}^Q \end{bmatrix} \tag{3}$$

where $\mathbf{X} \in \mathbb{N}_0^{C \times G}$ for $C = \sum_{q=1}^{Q} C_q$ cells and $G$ genes, and $\mathbf{X}^q \in \mathbb{N}_0^{C_q \times G}$ denotes the single-cell expression count matrices from the control ($q = 1$) and each treatment level ( $q = 2, \ldots, Q$).

#### The loading weight tensor **W**

In order to account for the conditioning variable, we introduce for each level a $F \times G$ gene loading weights, giving rise to a $F \times G \times Q$ sized loading tensor $\mathbf{W}$. Another way to conceptualize this is to view $\mathbf{W}$ as a stacked array of $Q$ loading matrices

$$\mathbf{W} = \left[\mathbf{W}^1; \mathbf{W}^2; \ldots; \mathbf{W}^Q\right] \tag{4}$$

where each $\mathbf{W}^q = \{w_{fgq}\}$ for $q = 1, \ldots, Q$ represents a $F \times G$ matrix.

#### The loading weight design matrix **D**

To inform the factorisation framework about the conditioning variable, we utilize full-column rank design matrices. For the expression count matrix in Eq (3), $\mathbf{D}$ takes the form

$$\mathbf{D} = \begin{bmatrix} \mathbf{1}_{C_1} & \mathbf{0}_{C_1} & \mathbf{0}_{C_1} & \ldots & \mathbf{0}_{C_1} \\ \mathbf{1}_{C_2} & \mathbf{1}_{C_2} & \mathbf{0}_{C_2} & \ldots & \mathbf{0}_{C_2} \\ \mathbf{1}_{C_3} & \mathbf{0}_{C_3} & \mathbf{1}_{C_3} & \ldots & \mathbf{0}_{C_3} \\ \vdots & \vdots & \vdots & \ddots & \vdots \\ \mathbf{1}_{C_Q} & \mathbf{0}_{C_Q} & \mathbf{0}_{C_Q} & \ldots & \mathbf{1}_{C_Q} \end{bmatrix} \in \mathbb{R}^{C \times Q} \tag{5}$$

where $\mathbf{1}_{C_q}$ and $\mathbf{0}_{C_q}$ represent column vectors of length $C_q$ for $q = 1, \ldots, Q$ with ones and zeros, respectively. Note that the first column of $\mathbf{D}$ encodes the reference level and contains only ones. All other columns $2, \ldots, Q$ contain ones in rows where $\mathbf{X}$ (Eq. (3)) harbors cells of the respective condition and otherwise zeros. Statistically inclined readers will recognise that the form of $\mathbf{D}$ is reminiscent of design matrices employed in multiple linear regression (MLR) models with indicator variables. Similar to MLR models, where the design matrices parameterise regression coefficients, scPCA leverages $\mathbf{D}$ to sum slices of the loading tensor $\mathbf{W}^q$ (see Eq. (4)) in a condition-dependent manner.

#### The mean offset matrix **V**

In scPCA, we directly model single-cell expression data in count space (Eq. (1)), thereby evading the need to transform the data prior to the analysis. However, in order to find the condition-specific bases it is mandatory to center the data[1]. For this reason, we introduce the parameter $\mathbf{V}$ which captures the mean expression of each gene.

#### The mean offset indicator matrix **B**

We enable parameterizing $\mathbf{V}$ by providing an optional indicator matrix $\mathbf{B}$. We found that a global mean offset suffices for most multi-condition datasets. In this case, $\mathbf{B}$ reduces to a column vector of ones. However, if the experimental intervention causes shifts in mean gene expression, distinct mean offsets for each condition are crucial to ensure alignment of the bases. Then, $\mathbf{B}$ takes the form of an indicator matrix

$$\mathbf{B} = \begin{bmatrix} \mathbf{1}_{C_1} & \mathbf{0}_{C_1} & \mathbf{0}_{C_1} & \ldots & \mathbf{0}_{C_1} \\ \mathbf{0}_{C_2} & \mathbf{1}_{C_2} & \mathbf{0}_{C_2} & \ldots & \mathbf{0}_{C_2} \\ \mathbf{0}_{C_3} & \mathbf{0}_{C_3} & \mathbf{1}_{C_3} & \ldots & \mathbf{0}_{C_3} \\ \vdots & \vdots & \vdots & \ddots & \vdots \\ \mathbf{0}_{C_Q} & \mathbf{0}_{C_Q} & \mathbf{0}_{C_Q} & \ldots & \mathbf{1}_{C_Q} \end{bmatrix}. \tag{6}$$

Finally, we compute a cell-specific size factor $l_c = \log \sum_{g=1}^{G} x_{cg}$ which accounts for technical differences in sequencing depth between cells.

### Model Parameterisation and Intuition

Consider a conditioning variable with $Q = 2$ levels. For example, we might be interested in jointly modeling single-cell data from a control (ctl) and treatment (trt) experiment comprising $C_1$ ctl and $C_2$ trt cells, respectively. The concatenated transcript count matrix $\mathbf{X} \in \mathbb{N}_0^{C \times G}$ where $C = C_1 + C_2$ may then take the form

$$\mathbf{X} = \begin{bmatrix} \mathbf{X}^{\text{ctl}} \\ \mathbf{X}^{\text{trt}} \end{bmatrix}$$

where $\mathbf{X}^{\text{ctl}} \in \mathbb{N}_0^{C_1 \times G}$ and $\mathbf{X}^{\text{trt}} \in \mathbb{N}_0^{C_2 \times G}$ contain the expression data from ctl and trt cells, respectively.

Choosing the ctl condition as reference, we consider the following design and indicator matrices

$$\mathbf{D} = \begin{bmatrix} \mathbf{1}_{C_1} & \mathbf{0}_{C_1} \\ \mathbf{1}_{C_2} & \mathbf{1}_{C_2} \end{bmatrix} \quad \text{and} \quad \mathbf{B} = \begin{bmatrix} \mathbf{1}_{C_1} & \mathbf{0}_{C_1} \\ \mathbf{0}_{C_2} & \mathbf{1}_{C_2} \end{bmatrix}$$

where $\mathbf{1}_{C_q}$ and $\mathbf{0}_q$ represent column vectors of length $C_q$ for $q = \{1, 2\}$ with ones and zeros, respectively. Denote $\mathbf{d}_c$ as the $c$th row vector of $\mathbf{D}$ (and similarly for all other matrices). Note that a cell $i$ from the ctl condition is encoded with the row vectors $\mathbf{d}_i = [1, 0]$ and $\mathbf{b}_i = [1, 0]$, respectively. Conversely, a cell $j$ from the trt condition is represented by row vectors in the form of $\mathbf{d}_j = [1, 1]$ and $\mathbf{b}_j = [0, 1]$, respectively.

To understand how the design and indicator matrices $\mathbf{D}$ and $\mathbf{B}$ parameterise the model, we will now focus on the factor model term $\mathbf{S}$ and intercept term $\mathbf{T}$ separately (compare Eq. (2)), i.e.

$$\log \mu_{cg} = l_c + \underbrace{\left( \sum_{p=1}^{P} b_{cp} v_{pg} \right)_{cg}}_{\mathbf{T}} + \underbrace{\left( \sum_{q=1}^{Q} d_{cq} \cdot \sum_{f=1}^{F} z_{cf} w_{fgq} \right)_{cg}}_{\mathbf{S}} . \tag{7}$$

The parameterisation of the loading tensor **W**

We start by expanding the term $\mathbf{S}$ of Eq. (7) for a particular row $c$

$$\mathbf{s}_c = d_{c1} \cdot \textstyle\sum_{f=1}^{F} z_{cf} w_{fg1} + d_{c2} \cdot \textstyle\sum_{f=1}^{F} z_{cf} w_{fg2}$$

and explicitly write out the case for a cell $c = i$ belonging to the control group (i.e. $\mathbf{d}_i = [1, 0]$)

$$\begin{aligned} \mathbf{s}_i &= 1 \cdot \sum_{f=1}^{F} z_{if} w_{fg1} + 0 \cdot \sum_{f=1}^{F} z_{if} w_{fg2} \\ &= \sum_{f=1}^{F} z_{if} w_{fg1} \\ &= \mathbf{z}_i \mathbf{W}^1 = \mathbf{z}_i \mathbf{W}^{\text{ctl}} \end{aligned}$$

and likewise for a treated cell $c = j$ (i.e. $\mathbf{d}_j = [1, 1]$ )

$$\begin{aligned} \mathbf{s}_j &= 1 \cdot \sum_{f=1}^{F} z_{jf} w_{fg1} + 1 \cdot \sum_{f=1}^{F} z_{jf} w_{fg2} \\ &= \sum_{f=1}^{F} z_{jf} \left( w_{fg1} + w_{fg2} \right) \\ &= \mathbf{z}_j \left( \mathbf{W}^1 + \mathbf{W}^2 \right) = \mathbf{z}_j \left( \mathbf{W}^{\text{ctl}} + \mathbf{W}^2 \right) = \mathbf{z}_j \mathbf{W}^{\text{trt}}. \end{aligned}$$

We find that the design matrix instructs the model to sum slices of the loading tensor in a cell/condition-dependent manner. Note that $\mathbf{D}$'s first column contains ones to designate the user-specified reference group. As a consequence, the bases for each other experimental condition encoded in $\mathbf{D}$ is a linear combination of the first slice $\mathbf{W}^1$ which represents the loading matrix of the ctl group ($\mathbf{W}^{\text{ctl}}$). We stress the importance of expressing the loading matrices of an experimental condition with respect to the specified reference because it allows the model to fit aligned components.

#### The parameterisation of the loading tensor V

We now focus on the intercept term of Eq. (7). We observed that supplying an indicator matrix is optional for most single-cell datasets, in which case the $\mathbf{B}$ reduces to a vector of ones $\mathbf{1}_C$ and global mean offsets are inferred. However, in cases when it is reasonable to assume that there are substantial deviations in mean expression across conditions, it may be mandatory to center gene expression in a condition-specific manner. Then $\mathbf{B}$ takes the form of an indicator matrix (see Eq. (6)).

We first expand term $\mathbf{T}$ from Eq. (7) for a particular row $c$

$$\mathbf{t}_c = b_{c1} \cdot v_{1g} + b_{c2} \cdot v_{2g}$$

and explicitly write out the case for a cell $c = i$ belonging to the ctl group (i.e. $\mathbf{b}_i = [1, 0]$), then

$$\begin{aligned} \mathbf{t}_i &= 1 \cdot v_{1g} + 0 \cdot v_{2g} \\ &= \mathbf{v}_1 = \mathbf{v}^{\text{ctl}} \end{aligned}$$

and likewise for a cell $c = j$ that is member of the trt group (i.e. $\mathbf{b}_j = [0, 1]$) reduces to

$$\begin{aligned} \mathbf{t}_j &= 0 \cdot v_{1g} + 1 \cdot v_{2g} \\ &= \mathbf{v}_2 = \mathbf{v}^{\text{trt}}. \end{aligned}$$

Taken together, we find that the intercept design matrix $\mathbf{B}$ effectively extracts distinct rows from $\mathbf{V}$ in a cell-specific manner, thereby allowing to model the mean across the $P$ groups defined in the indicator matrix $\mathbf{B}$ .

### Implementation and Inference

We express the scPCA in terms of a hierarchical Bayesian model. The generative model may be expressed as

$$
\begin{aligned}
x_{cg} \mid \mu_{cg}, \alpha_g &\sim \mathrm{NB}\left(\mu_{cg}, \alpha_g\right) \\
\log \mu_{cg} &= l_c + \left(\sum_{p=1}^{P} b_{cp} v_{pg}\right)_{cg} + \left(\sum_{q=1}^{Q} d_{cq} \cdot \sum_{f=1}^{F} z_{cf} w_{fgq}\right)_{cg} \\
z_{cf} &\sim \mathcal{N}(0, 0.1) \\
w_{fgq} &\sim \mathcal{N}(0, 1) \\
v_{pg} &\sim \mathcal{N}(0, 1) \\
\alpha_g^{-1} &\sim \mathrm{Exp}(\beta) \\
\beta &\sim \mathrm{Gamma}(3, 0.01).
\end{aligned} \tag{4}
$$

We choose Normal distributions as priors for the latent variables $\mathbf{W}$, $\mathbf{V}$ and $\mathbf{Z}$ of the model. Observe that we posit a scale of 0.1 on the prior for $\mathbf{Z}$ thereby encouraging the model to capture spectral differences in gene expression across conditions in the loading tensor $\mathbf{W}$ rather than to explain them with additional factors.

We use a containment prior to model the overdispersion $\alpha_g$ of each gene. Apriori the model assumes equidispersed (poissonian) counts by placing most probability mass on large $\alpha_g$. Note that we tightly fix the mean exponential rate $\beta$ around $3$ with a standard deviation of $0.01$.

Using matrix notation we may express the priors of the model as follows

$$
\begin{aligned}
p(\beta) &= \mathrm{Gamma}(\beta; 3, 0.01) \\
p\left(\boldsymbol{\alpha}^{-1} \mid \beta\right) &= \prod_{g=1}^{G} \mathrm{Exp}\left(\alpha_g^{-1} \mid \beta\right) \\
p(\mathbf{V}) &= \prod_{p=1}^{P} \prod_{g=1}^{G} \mathcal{N}\left(v_{pg}; 0, 1\right) = \prod_{p=1}^{P} \mathcal{N}\left(\mathbf{v}_p; \mathbf{0}_G, \mathbf{I}_G\right) \\
p(\mathbf{W}) &= \prod_{q=1}^{Q} \prod_{f=1}^{F} \prod_{g=1}^{G} \mathcal{N}\left(w_{fgq}; 0, 1\right) = \prod_{q=1}^{Q} \prod_{f=1}^{F} \mathcal{N}\left(\mathbf{w}_f^q; \mathbf{0}_G, \mathbf{I}_G\right) \\
p(\mathbf{Z}) &= \prod_{c=1}^{C} \prod_{f=1}^{F} \mathcal{N}\left(z_{cf}; 0, 0.1\right) = \prod_{c=1}^{C} \mathcal{N}\left(\mathbf{z}_c; \mathbf{0}_F, 0.1 \cdot \mathbf{I}_F\right)
\end{aligned}
$$

where $\mathbf{0}_A$ represents a $A$-dimensional vector of zeros and $\mathbf{I}_A$ an $A \times A$ identity matrix. Together with the likelihood of the data

$$
p\left(\mathbf{X} \mid \mathbf{W}, \mathbf{Z}, \mathbf{V}, \boldsymbol{\alpha}^{-1}\right) = \prod_{c=1}^{C} \prod_{g=1}^{G} \mathrm{NB}\left(x_{cg} \mid \mu_{cg}, \alpha_g\right)
$$

we derive the joint distribution

$$
\begin{aligned}
p\left(\mathbf{X}, \mathbf{W}, \mathbf{Z}, \mathbf{V}, \boldsymbol{\alpha}^{-1}, \beta\right) = & p(\beta) p\left(\boldsymbol{\alpha}^{-1} \mid \beta\right) p(\mathbf{W}) p(\mathbf{V}) p(\mathbf{Z}) p\left(\mathbf{X} \mid \mathbf{W}, \mathbf{Z}, \mathbf{V}, \boldsymbol{\alpha}^{-1}\right) \\
= & \operatorname{Gamma}(\beta ; 3,0.01) \\
& \times \prod_{p=1}^{P} \mathcal{N}\left(\mathbf{v}_p ; \mathbf{0}_G, \mathbf{I}_G\right) \\
& \times \prod_{q=1}^{Q} \prod_{f=1}^{F} \mathcal{N}\left(\mathbf{w}_f^q ; \mathbf{0}_G, \mathbf{I}_G\right) \\
& \times \prod_{g=1}^{G} \operatorname{Exp}\left(\alpha_g^{-1} ; \beta\right) \prod_{c=1}^{C} \operatorname{NB}\left(x_{cg} \mid \mu_{cg}, \alpha_g\right) \mathcal{N}\left(\mathbf{z}_c ; \mathbf{0}_F, 0.1 \cdot \mathbf{I}_F\right).
\end{aligned}
$$

We use stochastic variational inference in order the determine the posterior distribution of the latent variables of the model $p(\boldsymbol{\theta}, \boldsymbol{\alpha}^{-1}, \beta | \mathbf{X})$ where $\boldsymbol{\theta} = \{\mathbf{W}, \mathbf{V}, \mathbf{Z}\}$. Variational methods require us to define a surrogate posterior $q\left(\boldsymbol{\theta}, \boldsymbol{\alpha}^{-1}, \beta ; \boldsymbol{\phi}\right)$ which serves as an approximation to the true posterior. For this reason, we posit a mean-field approximation comprising normal and log normal distributions

$$
q\left(\boldsymbol{\theta}, \boldsymbol{\alpha}^{-1}, \beta ; \boldsymbol{\phi}\right) = \log \mathrm{N}\left(\beta ; m_1, s_1\right) \prod_{k=2}^{G+1} \log \mathrm{N}\left(\alpha_g^{-1} ; m_k, s_k\right) \prod_{k=G+2}^{K} \mathcal{N}\left(\theta_k ; m_k, s_k\right)
$$

where $\phi = (m_1, \ldots, m_K, s_1, \ldots, s_K)$ concatenates the mean and standard deviation of each Gaussian factor, and $K = 1 + G + P \times G + F \times G \times Q + C \times F$ represents the total number of parameters in the model.

We aim to find the variational parameters $\phi$ that minimize the KL divergence between surrogate and true posterior $\mathrm{KL}\left(q\left(\boldsymbol{\theta}, \boldsymbol{\alpha}^{-1}, \beta ; \boldsymbol{\phi}\right) \| p\left(\boldsymbol{\theta}, \boldsymbol{\alpha}^{-1}, \beta \mid \mathbf{X}\right)\right)$. However, since $p\left(\boldsymbol{\theta}, \boldsymbol{\alpha}^{-1}, \beta \mid \mathbf{X}\right)$ is unknown, we instead use the evidence lower bound (ELBO)[2]

$$
\mathcal{L}(\boldsymbol{\phi}) = \mathbb{E}_{q(\boldsymbol{\theta}, \boldsymbol{\alpha}^{-1}, \beta)}\left[\log p\left(\mathbf{X}, \boldsymbol{\theta}, \boldsymbol{\alpha}^{-1}, \beta\right) - \log q\left(\boldsymbol{\theta}, \boldsymbol{\alpha}^{-1}, \beta ; \boldsymbol{\phi}\right)\right]
$$

which is equivalent to the KL divergence up to a constant. We use a gradient descent-based method (black box variational inference) to solve the optimisation problem

$$
\phi^* = \arg\max_\phi \mathcal{L}(\phi)
$$

which requires us to estimate the gradients of the expectation in Eq. (5) with respect to the variational parameters $\phi$. In practice this may be achieved using Monte Carlo integration,

$$
\nabla_{\boldsymbol{\phi}} \mathcal{L}(\boldsymbol{\phi}) \approx \sum_{t=1}^{T} \nabla_{\boldsymbol{\phi}}\left[\log p\left(\mathbf{X}, \boldsymbol{\theta}_t, \boldsymbol{\alpha}_t^{-1}, \beta_t\right) - \log q\left(\boldsymbol{\theta}_t, \boldsymbol{\alpha}_t^{-1}, \beta_t ; \boldsymbol{\phi}\right)\right]
$$

where $T$ corresponds to the number of samples we draw from $q\left(\boldsymbol{\theta}, \boldsymbol{\alpha}^{-1}, \beta ; \phi\right)$ during a single training step (by default $T = 1$).

We implemented scPCA using the pyro probabilistic programming language. By default we train the models for 30000 epochs, employ an ADAMGrad optimizer with a multi-step learning rate decay and an initial learning rate of 0.1. For scalability, we compute stochastic gradients over batches of observations.

### Relationship to other factor model approaches

In the following section we compare scPCA to other factor models and related modeling approaches which incorporate secondary variables into their analyses.

#### Principal component analysis (PCA)

PCA projects high-dimensional data into a lower-dimensional space, maximizing the variance explained by the projection. Consider a mean-centered data matrix $\mathbf{X} \in \mathbb{R}^{C \times G}$ for $C$ observations and $G$ features. To perform a PCA on $\mathbf{X}$ we may employ a singular value decomposition (SVD)

$$\mathbf{X} = \mathbf{U}\mathbf{S}\mathbf{V}^T \text{ where } \mathbf{U}^T\mathbf{U} = \mathbf{V}^T\mathbf{V} = \mathbf{I}$$

where $\mathbf{U}$ and $\mathbf{V}$ denote the left and right singular matrices, respectively, and $\mathbf{S}$ is a diagonal matrix containing the singular values. The decomposition reveals that $\mathbf{Z} = \mathbf{U}\mathbf{S}$ and $\mathbf{W} = \mathbf{V}^T$ such that

$$\mathbf{X} = \mathbf{Z}\mathbf{W}$$

where $\mathbf{Z}$ denotes the factor matrix and $\mathbf{W}$ the corresponding loading matrix. The linearity assumption ensures that the resulting loading vectors from PCA are simple and interpretable.

#### Probabilistic principal component analysis (PPCA)

The probabilistic formulation of PCA may be derived from a Gaussian latent variable model, which seeks to relate the $G$ -dimensional observations $\mathbf{X} = \{\mathbf{x}_c\}$ to a $F$-dimensional ( $F < G$ ) vector of latent variables through a linear model [3,4]. This relationship may be concisely denoted as

$$\mathbf{X} = \mathbf{Z}\mathbf{W} + \mathbf{E}$$

where $\mathbf{Z}$ a $C \times F$ factor matrix, $\mathbf{W}$ represents a $F \times G$-matrix loading matrix, and $\mathbf{E}$ a $C \times G$ residual matrix. Specifying an isotropic noise model induces a Gaussian on the marginal distribution of $\mathbf{x}$, i.e. $\mathbf{x} \sim \mathcal{N}(\boldsymbol{\mu}, \mathbf{W}\mathbf{W}^T + \sigma^2\mathbf{I})$, and it can be shown that as the covariance of the noise model becomes infinitesimal small $\sigma^2 \to 0$ standard PCA is recovered [3]. In this case, the $F$ rows of $\mathbf{W}$ span a linear subspace that corresponds to the principal subspace of the data.

There are many variations and extensions of the PPCA model that enable it to handle binary[5] or categorical data[6]. For example, contrastive PCA[7,8] identifies components by contrasting the variance in a target dataset against a background dataset, thus highlighting unique patterns specific to the target. The GLM PCA framework[1] introduced an expectation-maximization algorithm to facilitate modeling count data with an over-dispersed negative binomial distribution.

The scPCA model is closely related to PPCA but enables to account for conditioning variables. Specifically, it assumes that the principal subspaces for each condition are closely related, allowing the bases for each condition to be expressed as a linear transformation of a reference basis/condition.

#### Principal component regression (PCR)

In principal component regression (PCR), we aim to predict a dependent variable $\mathbf{Y}$ using high-dimensional data $\mathbf{X}$[9,10]. This is straightforward when $\mathbf{X}$ is invertible, but it becomes challenging in high-dimensional settings where $\mathbf{X}$ is singular due to multicollinearity. PCR addresses this by performing PCA on $\mathbf{X}$ and then using the resulting factors $\mathbf{Z}$ as independent variables in a multiple regression on $\mathbf{Y}$. Since the PCA factors are uncorrelated, this approach mitigates the multicollinearity problem that often complicates high-dimensional analysis.

However, PCR does not inherently identify an optimal subset of predictors in $\mathbf{X}$. Instead, inspecting the prominent loading weights of the factors with the largest regression coefficients may hint at important features. Also note that the factors are chosen to optimally explain $\mathbf{X}$ rather than $\mathbf{Y}$, meaning PCR does not guarantee find factors that represent meaningful (i.e. predictive) covariates for $\mathbf{Y}$.

#### Partial least squares (PLS)

Partial least squares (PLS) finds factors from $\mathbf{X}$ that best predict $\mathbf{Y}$. Unlike PCR, which builds on factors explaining the variance in $\mathbf{X}$, PLS aims to find a basis for $\mathbf{X}$ that maximizes the variance explained in $\mathbf{Y}$. To achieve this, PLS finds factors $\mathbf{Z}$ that enable the simultaneous decomposition of $\mathbf{X}$ and $\mathbf{Y}$ such that the components explain as much as possible of the covariance between $\mathbf{X}$ and $\mathbf{Y}$. Specifically, we require that

$$\mathbf{X} = \mathbf{Z}\mathbf{W}^1 \text{ with } \mathbf{Z}^T\mathbf{Z} = \mathbf{I}$$

and

$$\hat{\mathbf{Y}} = \mathbf{Z}\mathbf{S}\mathbf{W}^2$$

where $\mathbf{W}^1$ represents a loading weight matrix for $\mathbf{X}$, $\mathbf{S}$ is a diagonal matrix with scaling factors, and $\mathbf{W}^2$ may be considered as a “weight matrix” of the dependent variable $\mathbf{Y}$[9,11]. Several algorithms have been proposed to perform PLS [12–14].

#### Canonical Correlation analysis (CCA)

Canonical Correlation Analysis (CCA) is a classical method for identifying correlations between two multivariate data sets, $\mathbf{X}^1 \in \mathbb{R}^{N \times D_1}$ and $\mathbf{X}^2 \in \mathbf{R}^{N \times D_2}$ [15–17] for $N$ observations and $D_m$ features ($m = \{1, 2\}$). Specifically, CCA finds two bases for the data such that the correlation between the projections of these variables onto these bases is maximized. In other words, CCA determines linear transformations for $\mathbf{X}^1$ and $\mathbf{X}^2$ so that the transformed coordinates are maximally correlated. CCA shares some similarities with PLS regression, with the key difference being that CCA treats both $\mathbf{X}^1$ and $\mathbf{X}^2$ symmetrically, while PLS uses one of the datasets to predict the other. Klami et al. proposed a Bayesian CCA method that incorporates sparse priors on the loading weights to enhance interpretability [18]. This model may be expressed

$$\mathbf{X}^1 = \mathbf{Z}\mathbf{W}^1 + \mathbf{E}^1 \text{ and } \mathbf{X}^2 = \mathbf{Z}\mathbf{W}^2 + \mathbf{E}^2$$

where $\mathbf{W}^1 \in \mathbb{R}^{F \times D_1}$ and $\mathbf{W}^2 \in \mathbb{R}^{F \times D_2}$ represent loading weight matrices comprising $F$ factors, $\mathbf{Z} \in \mathbb{R}^{N \times F}$ a shared factor matrix and $\mathbf{E}^m \in \mathbb{R}^{N \times D_m}$ for $m = \{1, 2\}$ error matrices. Building on this, Multi-Omics Factor Analysis (MOFA)[19,20] generalized Bayesian CCA to accommodate multiple datasets. MOFA supports a range of likelihoods and incorporates sparsity priors, providing a more flexible and interpretable framework for analyzing complex data sets.

#### Surrogate variable analysis (SVA)

SVA is a factor model based approach originally developed for DNA microarray chips with the aim to enable the unbiased detection of differential expressed genes with respect to some conditioning variables [21,22]. Specifically, the authors attribute gene expression changes to the presence of "primary variables" (in this manuscript "conditioning variable") and other unmodeled variables (expression heterogeneity (EH)), which may not be studied explicitly because measurements are not available, or it is simply inconvenient to do so. For example, one may be interested in quantifying changes in expression due to a disease condition. However, the expression of certain genes may also depend on the age, and some genes may be affected by both disease condition and age. In this example, disease condition may be considered as a primary variable and age as a variable that contributes to expression heterogeneity. To account for EH the authors propose to

$$\mathbf{X} = \boldsymbol{\mu} + f(\mathbf{C}) + \mathbf{Z}\mathbf{W} + \mathbf{E}$$

where $\boldsymbol{\mu}$ represents a mean offset vector, $f(\mathbf{C})$ the effect of the primary variables, $\mathbf{Z}\mathbf{W}$ models EH, and $\mathbf{E}$ a residual matrix. To estimate $\mathbf{Z}\mathbf{W}$, the authors propose an iterative procedure in which they first subtract the signal due to the primary variables, and apply a low-rank decomposition to the resulting residual matrix. They then identify surrogate variables by pinpointing the subset of genes driving the signature of each significant EH component. Finally, the authors employ the surrogate variables as covariates for a regression analysis, thereby allowing to account for EH in the analysis.

In summary, while scPCA, (P)PCA and CCA reconstruct linear hyperplanes which encompass both primary and unmodelled variables, SVA explicitly tries to reconstruct a subspace that mainly explains expression heterogeneity. In this way, SVA enables an unbiased assessment of the effects of the primary variable on gene expression.

#### Remove Unwanted Variation (RUV-2)

The RUV-2 model was originally developed for microarray data[23], and may be considered as an extension to SVA. The authors propose to decompose the gene expression matrix

$$\mathbf{X} = \mathbf{C}\mathbf{W}^1 + \mathbf{Z}\mathbf{W}^2 + \mathbf{E}$$

where $\mathbf{C}$ is a matrix whose columns encode the conditioning variable of interest, $\mathbf{Z}$ represents unobserved (hidden) factors representative for unwanted variation, and $\mathbf{W}^1$ and $\mathbf{W}^2$ matrices that harbor coefficients measuring the influence of the conditioning variable or factor on a gene (analogously to loading matrices). Note that we have dropped here an optional term for known observed covariates which are not of interest (e.g. batch).

A major difference between SVA and RUV-2 is in the way that both algorithms estimate the factors of unwanted variation. While SVA first subtracts the signal due to the conditioning variable from the data and decomposes the residuals to estimate the subspace of unwanted variation, RUV leverages negative control genes, i.e. a set of genes which are known to be unassociated with the conditioning variable. In other words, to estimate $\mathbf{Z}$, the RUV-2 algorithm first decomposes

$$\mathbf{X}_c = \mathbf{Z}\mathbf{W}_c^2 + \mathbf{E}_c$$

where $c$ denotes the set of negative control genes and $\mathbf{W}_c^2$ a loading matrix, and $\mathbf{E}_c$ a residual matrix. The estimate of $\mathbf{Z}$ is then plugged into equation above and $\mathbf{W}^1$ and $\mathbf{W}^2$ are estimated via regression. The original RUV-2 model has been adapted to RNA-seq data[24,25].

#### Gene Expression Decomposition and Integration (GEDI)

The GEDI factorisation framework[26] prescribes a specific parametrization to model the lower-dimensional subspaces of multi-condition single cell data. Particularly, the authors propose

$$\mathbf{X} = \boldsymbol{\mu}_r + \boldsymbol{\mu}_{i(c)} + (\mathbf{W}_r + \mathbf{W}_{i(c)})\mathbf{Z} + \mathbf{E}$$

where $\boldsymbol{\mu}_r$ represents the origin point for the reference plane (mean offset) spanned by the basis defined in $\mathbf{W}_r$ , $\boldsymbol{\mu}_{i(c)}$ a sample-specific offset to the origin point of the reference ( $i(c)$ denotes an indicator function for cell $c$ ) and $\mathbf{W}_{i(c)}$ a sample-specific transformation of the reference basis, $\mathbf{Z}$ a factor matrix, and $\mathbf{E}$ a residual matrix.

In scPCA, we generalize the GEDI model by introducing design (and/or indicator) matrices to flexibly parameterise the mean offset vector and the loading matrices. This allows scPCA to model arbitrary experimental designs including multiple conditioning variables as well as interaction effects. Additionally, we model the single-cell expression count matrix as negative binomial distributed thus mitigating the need to preprocess the data.
