## Extended Data Figures for "Joint Modeling of Cellular Heterogeneity and Condition Effects with scPCA in Single-Cell RNA-Seq"

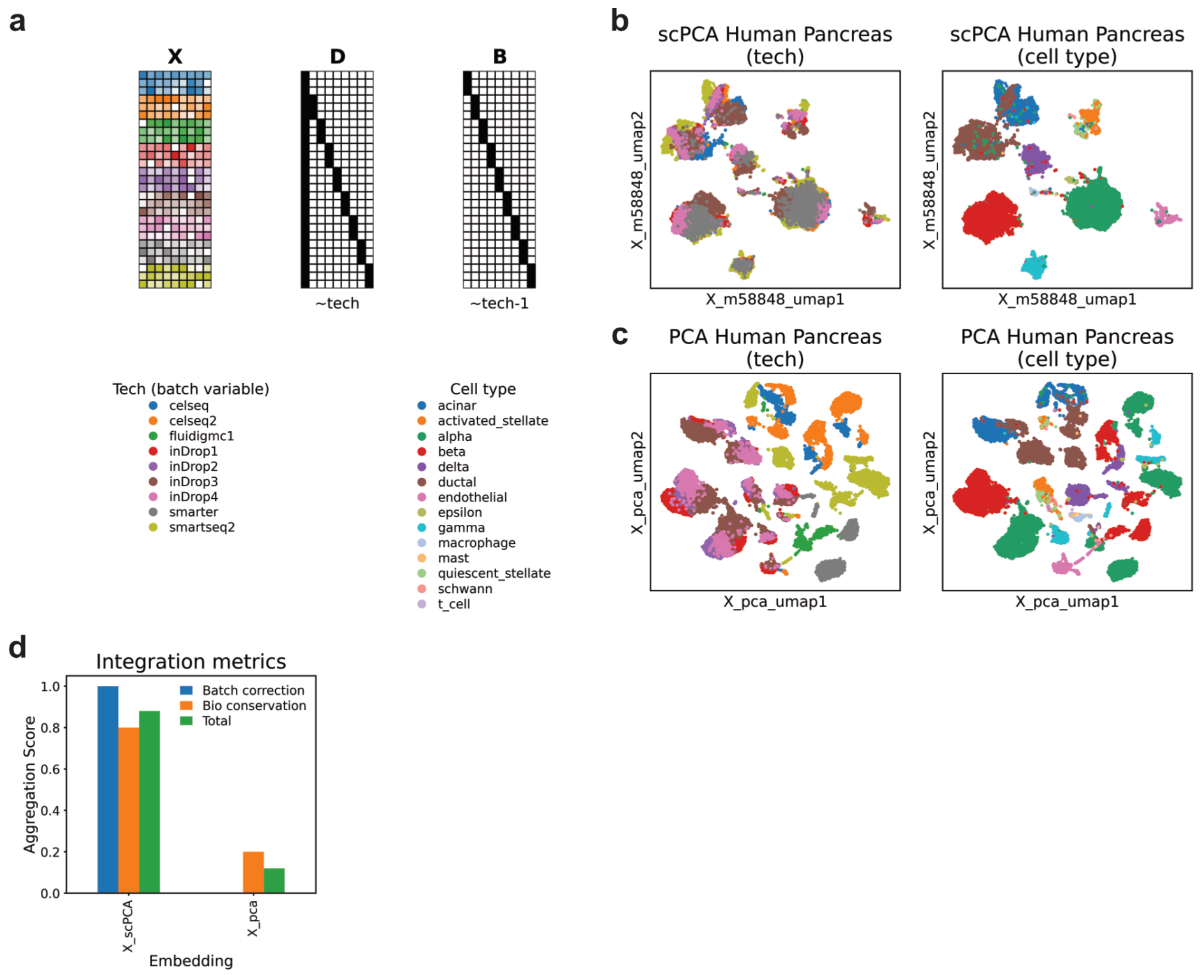


**Extended Data Fig. 1: Integrating the human pancreas single-cell dataset with scPCA.** **a.** Schematic depiction of the human pancreas dataset **X** comprising single-cell data from 9 different sequencing platforms, along with the corresponding design and indicator matrices **D** and **B**, which reflect the batch structure in the data. **b.** UMAP plots based on scPCA cell embeddings showing annotations for the batch variable (tech) and cell types. **c.** UMAP plots based on PCA cell embeddings showing annotations for the batch variable and cell types. **d.** Bio conservation and batch integration scores for PCA and scPCA embeddings.

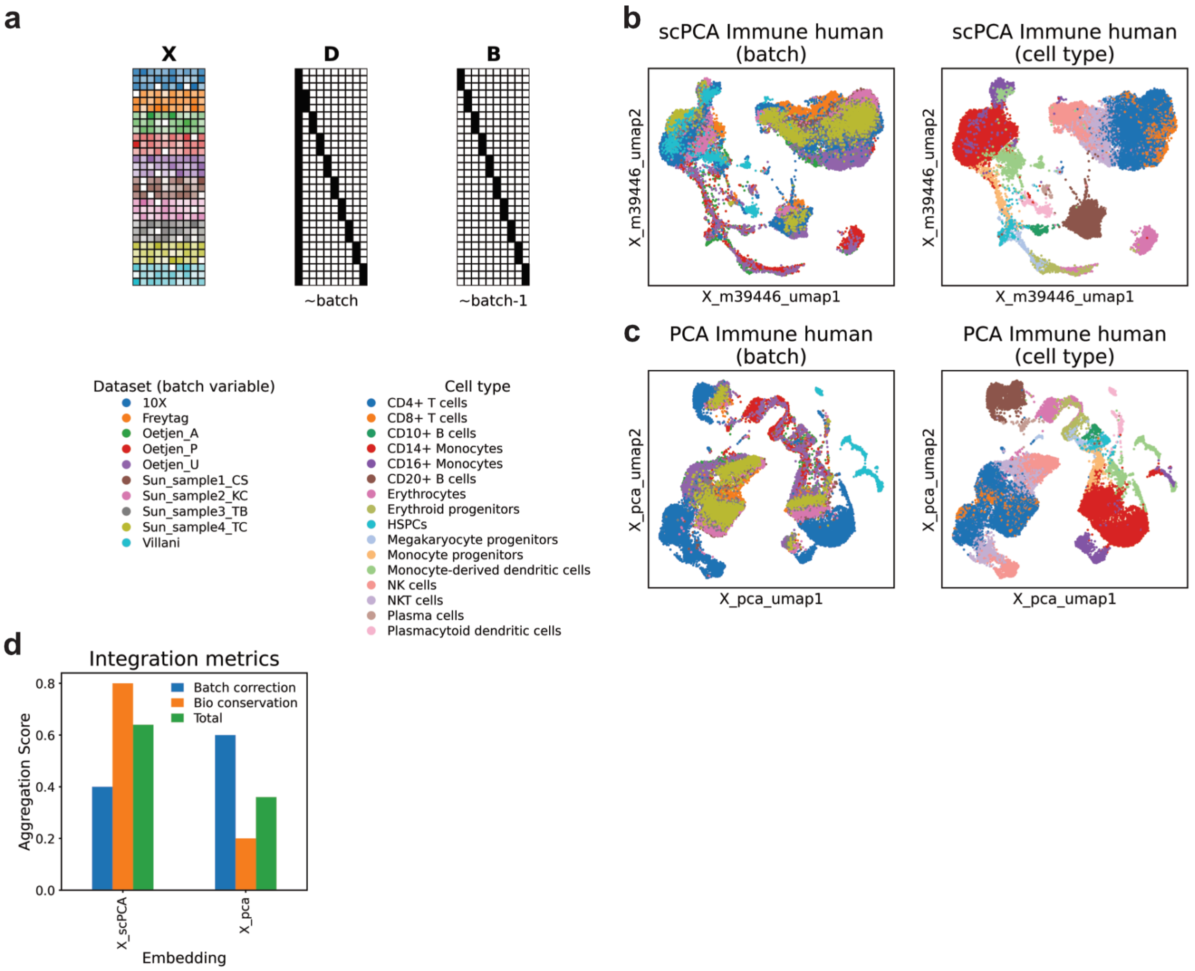


**Extended Data Fig. 2: Integrating the human immune dataset with scPCA. a.** Schematic depiction of the human immune dataset **X** comprising single-cell data from 10 donors, along with the corresponding design and indicator matrices **D** and **B**, which reflect the donor structure in the data. **b.** UMAP plots based on scPCA cell embeddings showing annotations for the batch variable and cell types. **c.** UMAP plots based on PCA cell embeddings showing annotations for the batch variable and cell types. **d.** Bio conservation and batch integration scores for PCA and scPCA embeddings.

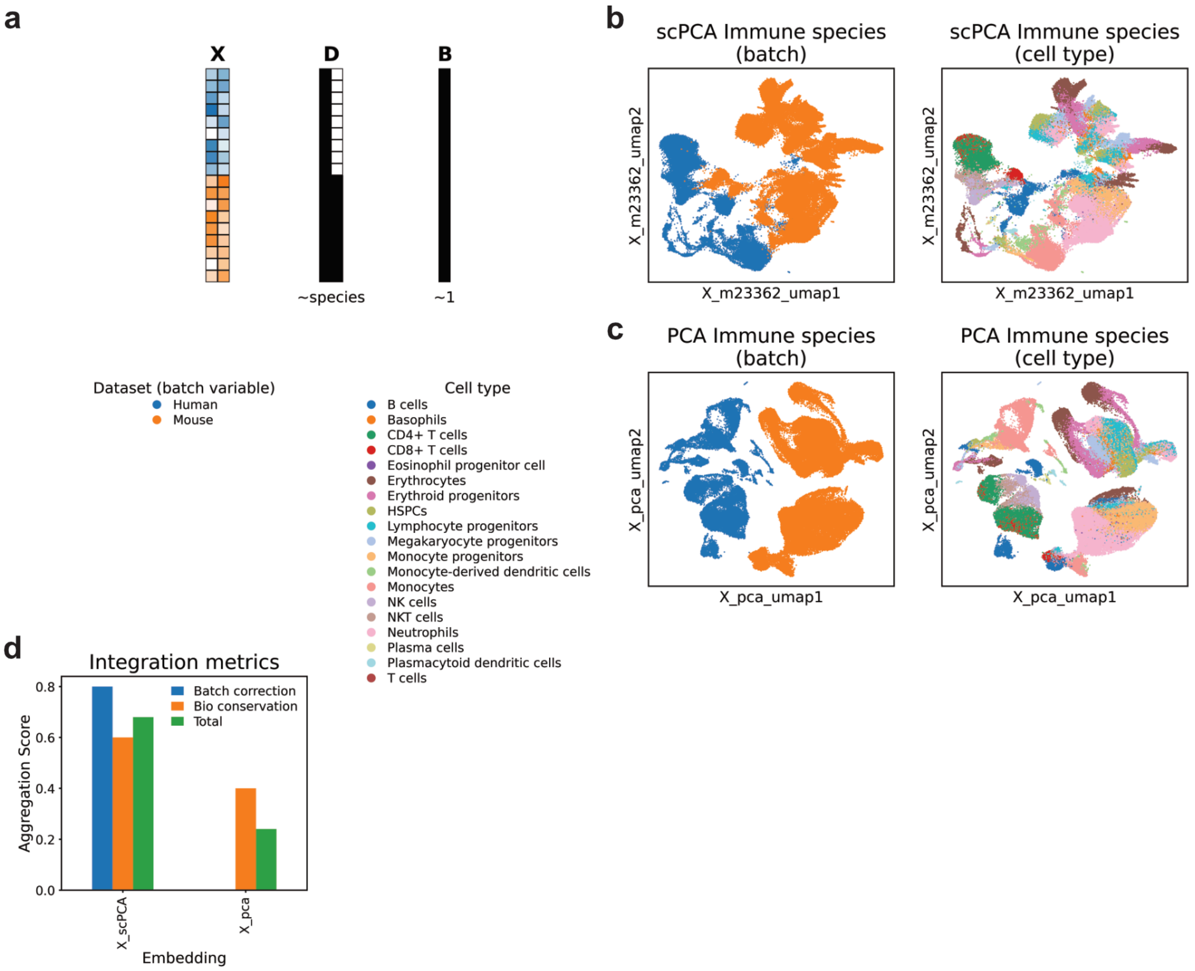


**Extended Data Fig. 3: Integrating immune cells across species using scPCA. a.** Schematic depiction of the mouse immune dataset **X** comprising single-cell data from 2 species, along with corresponding design and indicator matrices **D** and **B**, which reflect the batch structure in the data. **b.** UMAP plots based on scPCA cell embeddings showing annotations for the batch variable and cell types. **c.** UMAP plots based on PCA cell embeddings showing annotations for the batch variable and cell types. **d.** Bio conservation and batch integration scores for PCA and scPCA embeddings.

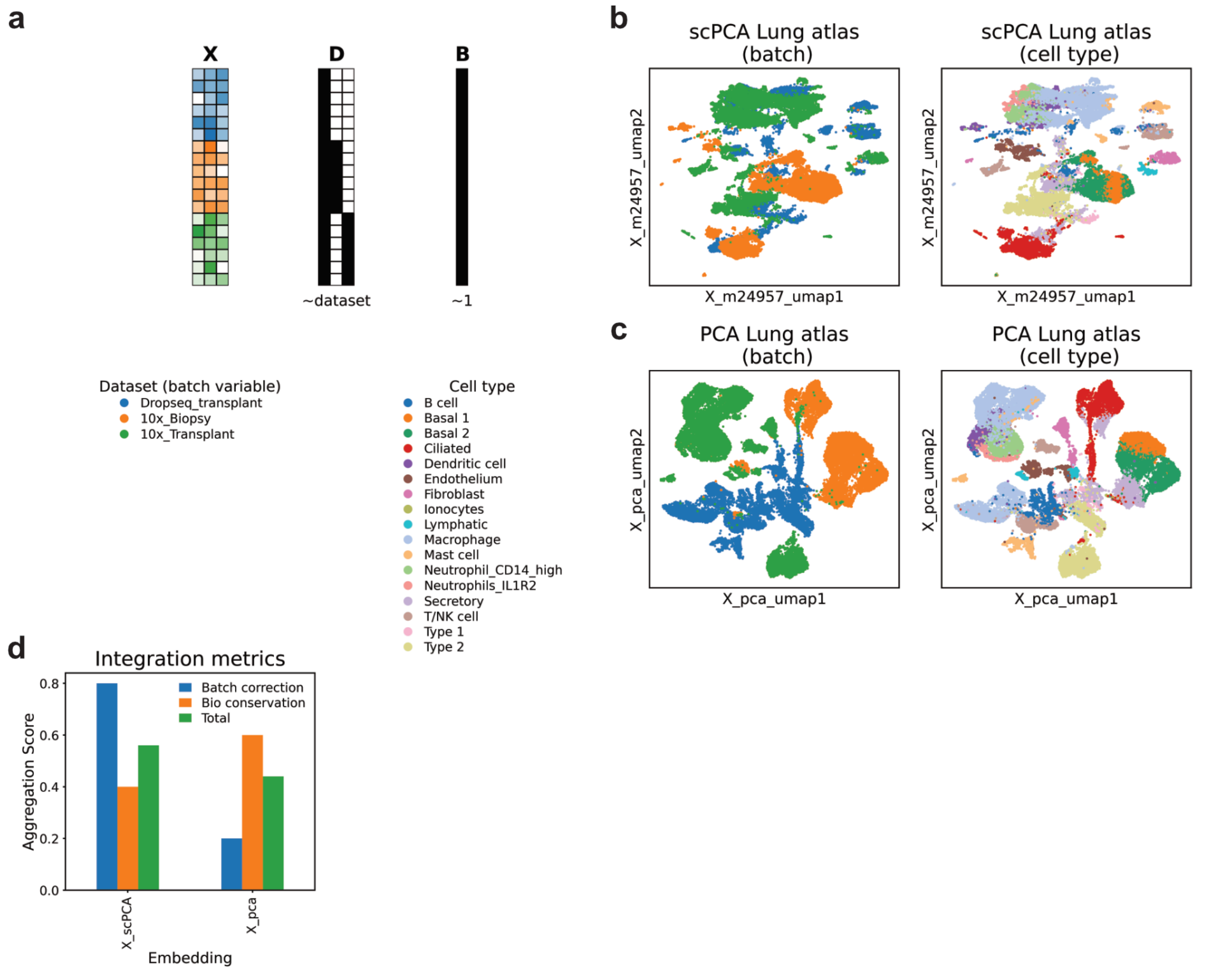


**Extended Data Fig. 4: Integrating the lung dataset across protocols with scPCA. a.** Schematic depiction of the lung dataset **X**, and corresponding design and indicator matrices **D** and **B**, which reflect the batch structure in the data. **b.** UMAP plots based on scPCA cell embeddings showing annotations for the batch variable and cell types. **c.** UMAP plots based on PCA cell embeddings showing annotations for the batch variable and cell types. **d.** Bio conservation and batch integration scores for PCA and scPCA embeddings.

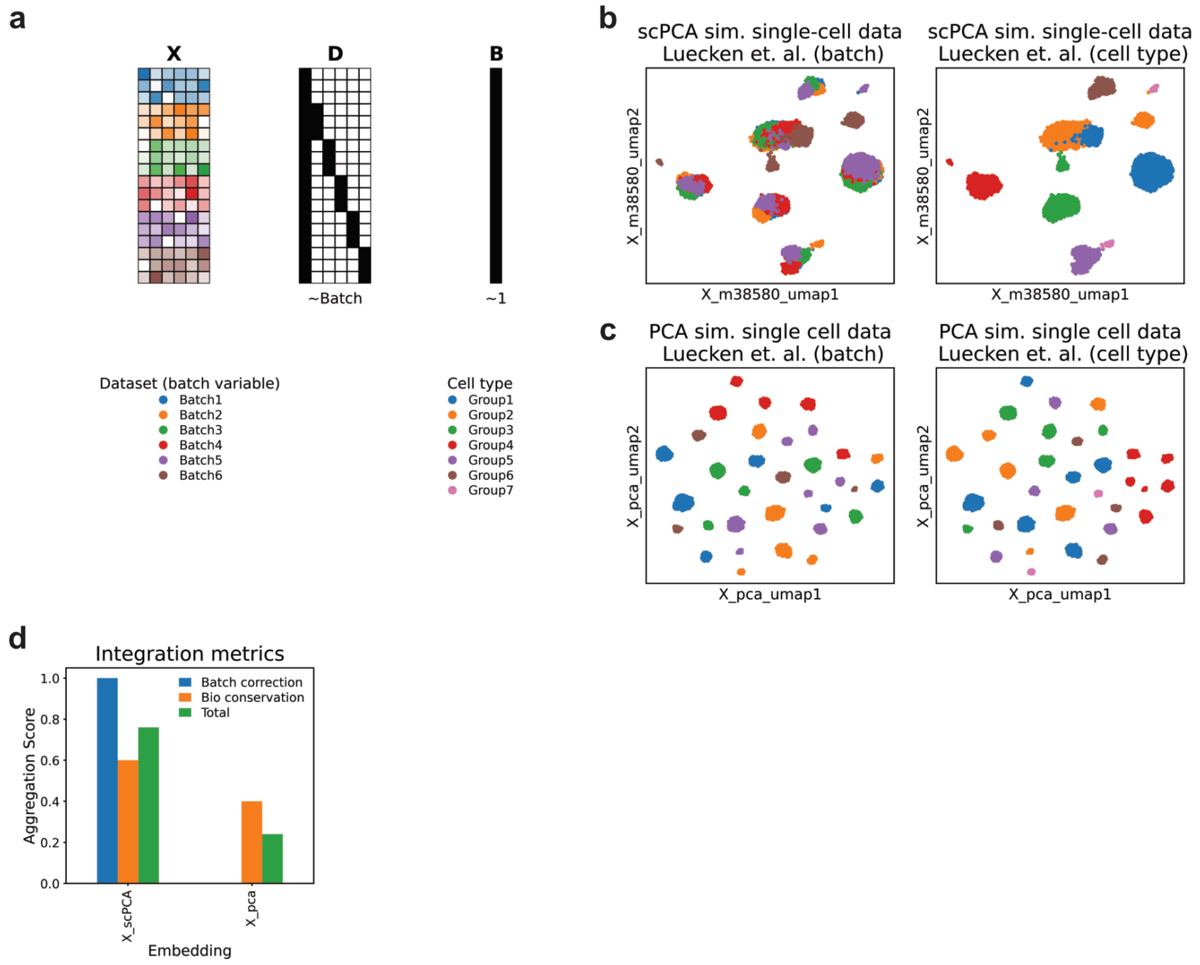


**Extended Data Fig. 5: Integrating simulated single-cell data with scPCA. a.** Schematic depiction of the simulated single-cell dataset by Luecken et al., and corresponding design and indicator matrices **D** and **B**, which reflect the batch structure in the data. **b.** UMAP plots based on scPCA cell embeddings showing annotations for the batch variable and cell types. **c.** UMAP plots based on PCA cell embeddings showing annotations for the batch variable and cell types. **d.** Bio conservation and batch integration scores for PCA and scPCA embeddings.

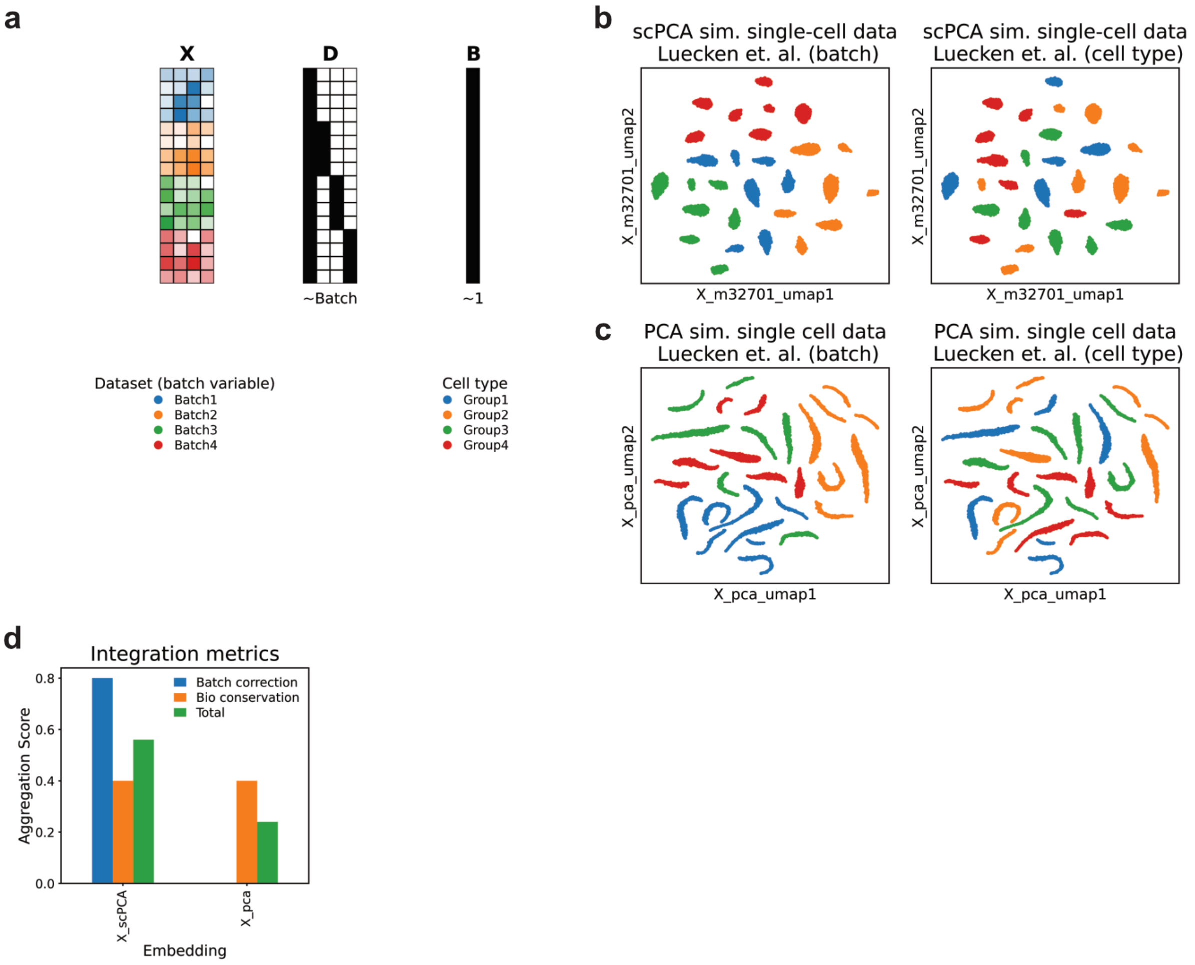


**Extended Data Fig. 6: Integrating complex simulated single-cell data with scPCA. a.** Schematic depiction of Luecken et al. second simulated single-cell dataset, and corresponding design and indicator matrices **D** and **B**, which reflect the batch structure in the data. **b.** UMAP plots based on scPCA cell embeddings showing annotations for the batch variable (tech) and cell types. **c.** UMAP plots based on PCA cell embeddings showing annotations for the batch variable and cell types. **d.** Bio conservation and batch integration scores for PCA and scPCA embeddings.
